# CD19×CD3 bispecific T cell engager treatment induces remission in experimental pemphigoid disease

**DOI:** 10.1101/2025.09.09.675030

**Authors:** Natalie Gross, Alexander Jochimsen, Sören Dräger, Thomas Theocharis, Andreas Recke, Nina van Beek, Leon Schmidt-Jiménez, Steffen Krohn, Katja Klausz, Katja Bieber, Ralf J. Ludwig, Matthias Peipp

## Abstract

Pemphigoid diseases (PDs) are autoimmune disorders marked by autoantibodies (Aab) against skin and mucous membrane proteins, causing muco-cutaneous blistering in predominantly elderly patients. Since current therapeutics are often insufficient, PDs impose a significant morbidity and mortality burden. CD19×CD3 bispecific T cell engagers (TCEs) - originally developed for B cell malignancies - have shown promise in treatment-refractory autoimmune diseases like systemic sclerosis and rheumatoid arthritis. To explore their potential in PDs, we tested a murine CD19×CD3 TCE in a mouse model of epidermolysis bullosa acquisita (EBA), a prototypical PD. The TCE selectively depleted B cells in blood and bone marrow, but not spleen, of healthy mice. Mice with clinically manifest immunization-induced EBA were randomized to receive either the CD19×CD3 TCE or control treatment upon reaching a predefined disease severity. After 13 weeks, 45% of TCE-treated mice achieved remission, versus 23% of controls. This improvement correlated with reduced antigen-specific B cells, though total and antigen-specific IgG levels were unchanged. These findings suggest that CD19×CD3 TCE preferentially target autoreactive B cells and may be effective at lower doses than those used in oncology. Our data support the potential of CD19×CD3 bispecific T cell engagers as a therapeutic strategy for PDs.

**eTOC Synopsis:** Gross, Jochimsen, Dräger and colleagues demonstrate that CD19×CD3 bispecific T cell engagers, can selectively target autoreactive B cells and induce remission in a mouse model of pemphigoid disease, highlighting their potential as a novel therapeutic strategy for autoimmune blistering disorders.

## Introduction

Autoimmune diseases pose a substantial medical burden. In many autoimmune diseases, pathogenic autoantibodies (Aab) have been identified. This makes Aab - and consequently Aab producing cells like B and plasma cells - promising therapeutic targets. Accordingly, treatments such as immunoadsorption, high doses of intravenous immunoglobulin G (IVIG), and rituximab have been established, either by directly removing circulating antibodies, promoting their degradation via the neonatal Fc receptor (FcRn), or depleting antibody-producing B cells.^1–3^ However, these therapies do not achieve consistent remission rates across all Aab-mediated diseases. Pemphigoid diseases (PDs), a group of chronic autoimmune skin blistering diseases, are one such example.^4^

In PDs, Aab targeting antigens of the dermal-epidermal junction (DEJ) drive disease pathology by recruiting effector cells such as neutrophils, T cells, and monocytes, ultimately leading to inflammation and subepidermal blistering.^5,6^ Bullous pemphigoid, the most common subtype of PD, predominantly affects elderly individuals with an average age between 78 to 80 years, depending on the study population.^7–9^ Given that nonspecific immunosuppression, such as systemic corticosteroids, substantially increases morbidity and mortality in this vulnerable patient population, there remains a critical unmet medical need for safer and more effective therapies. This is underscored by the limited efficacy of the CD20-targeting antibody rituximab.^10^ This highlights the need for more targeted approaches that effectively deplete autoreactive B cells while minimizing systemic immunosuppression.^4,11–13^

Two emerging strategies specifically targeting B cells are chimeric antigen receptor T (CAR-T) cells and bispecific antibodies. Both have been recently reviewed elsewhere.^14^ The efficacy of CAR-T or CAAR-T cells has been demonstrated in experimental models and patients, such as DSG3-CAAR-T cells in mucosal pemphigus vulgaris.^15–17^ However, high production costs,^18^ a complex manufacturing process, the need for pre-infusion conditioning with cell-depleting therapy, and - although minimal - the risk of CAR-T cell lymphomas all constrain their broad clinical application,^19^ particularly in elderly and multimorbid patients. Bispecific antibodies, such as blinatumomab, have emerged as potential therapeutics for autoimmune diseases.^20–24^ While conventional therapeutic IgG1 antibodies bind to a single antigen and predominantly mediate target cell killing via Fc-mediated effector functions like complement activation, antibody-dependent cell-mediated cytotoxicity (ADCC) and antibody-dependent cellular phagocytosis (ADCP), bispecific antibodies are engineered to simultaneously recognize two distinct antigens. In the case of bispecific T cell engaging antibodies, one domain binds a target cell associated antigen, whereas the other recruit T cells, typically by binding to CD3. This interaction facilitates the formation of an immunological synapse, leading to T cell activation and potent target cell lysis. Numerous bispecific antibody formats have been developed, and some have received regulatory approval for clinical use, especially in hematological malignancies. Examples include blinatumomab (CD19×CD3, for relapsed/refractory B-cell precursor acute lymphocytic leukemia) and teclistamab (BCMA**×**CD3, for relapsed/refractory multiple myeloma).^25,26^ Compared to CAR-T therapy, bispecific T cell engagers (TCEs) offer the advantage of being immediately available off-the-shelf. Both therapies can cause side effects like cytokine release syndrome (CRS) and neurotoxicity (ICANS). However, progress in biomarkers, dosing strategies, and safer constructs is improving their safety and efficacy.^27^ However, since patients with PD are often multimorbid,^9,12,13,28,29^ and therefore particularly vulnerable, thorough preclinical evaluation is essential before advancing new therapies to clinical use. To this end, we used the established immunization-induced mouse model of epidermolysis bullosa acquisita (EBA),^30,31^ which enables the investigation of therapeutic interventions in a controlled experimental setting. This model represents an ideal pre-clinical model to assess the efficacy of our CD19×CD3 TCE construct. In this model, mice are immunized with murine type VII collagen (COL7), leading to pathogenic Aab production and EBA development. The model has been extensively validated and widely used to study PD pathogenesis and evaluate potential therapeutics.^5^

To explore the concept of specifically targeting B cells in PD, we generated a murine CD19×CD3 bispecific antibody, validated its B cell-depleting activity in healthy mice, and ultimately evaluated its therapeutic efficacy in the immunization-induced EBA model.

## Results

### Design and in vitro Characterization of CD19×CD3 TCE

To develop the bispecific CD19×CD3, a TCE format was utilized in which a Fab fragment was coupled with a scFv through a flexible linker. Here, the Fab fragment of the murine CD3 antibody (clone 145-2C11) was linked to the VL and VH regions of a murine CD19 antibody (clone 1D3) via glycine-serine linkers (**Figure 1A**). The TCE was produced in CHO-S cells via transient transfection and subsequently purified through affinity chromatography followed by size-exclusion chromatography (SEC). Re-analysis of the purified molecule using SEC revealed a major peak at the expected retention volume (**Figure 1B**). Furthermore, SDS-PAGE under reducing and non-reducing conditions, followed by Coomassie blue staining, confirmed purity. As anticipated, two bands corresponding to the CD3 light chain (25 kDa) and the heavy chain derivative (55 kDa) were observed at the calculated molecular masses under reducing conditions. Under non-reducing conditions, a band for the assembled antibody was detected at a molecular mass of 75 kDa (**Figure 1C**). To confirm binding specificity, murine CD19^+^ B cells and CD3^+^ T cells were isolated from the spleen and incubated with varying concentrations of the antibody. The CD19×CD3 TCE exhibited comparable affinities for CD19 and CD3, with half-maximal effective concentrations (EC_50_) of 19.5 nM and 16.46 nM, respectively. Differences in maximum MFI values at saturation were attributed to varying antigen densities on the cell surface (**Figure 1D-E**). To evaluate the ability of the CD19×CD3 TCE to mediate T cell-dependent killing of CD19 cells, chromium release assays were performed using CHO-S cells transiently expressing murine CD19 as target cells. The TCE demonstrated efficient target cell lysis in a concentration-dependent manner, with an EC_50_ value of 12.69 pM. In contrast, the control antibody (Ctrl**×**CD3) showed no target cell lysis (**Figure 1F**). In conclusion, the CD19×CD3 TCE exhibited high-affinity binding to CD19^+^ target cells and CD3^+^ effector cells and mediated potent target cell lysis in vitro at picomolar concentrations.

**Figure 1:**
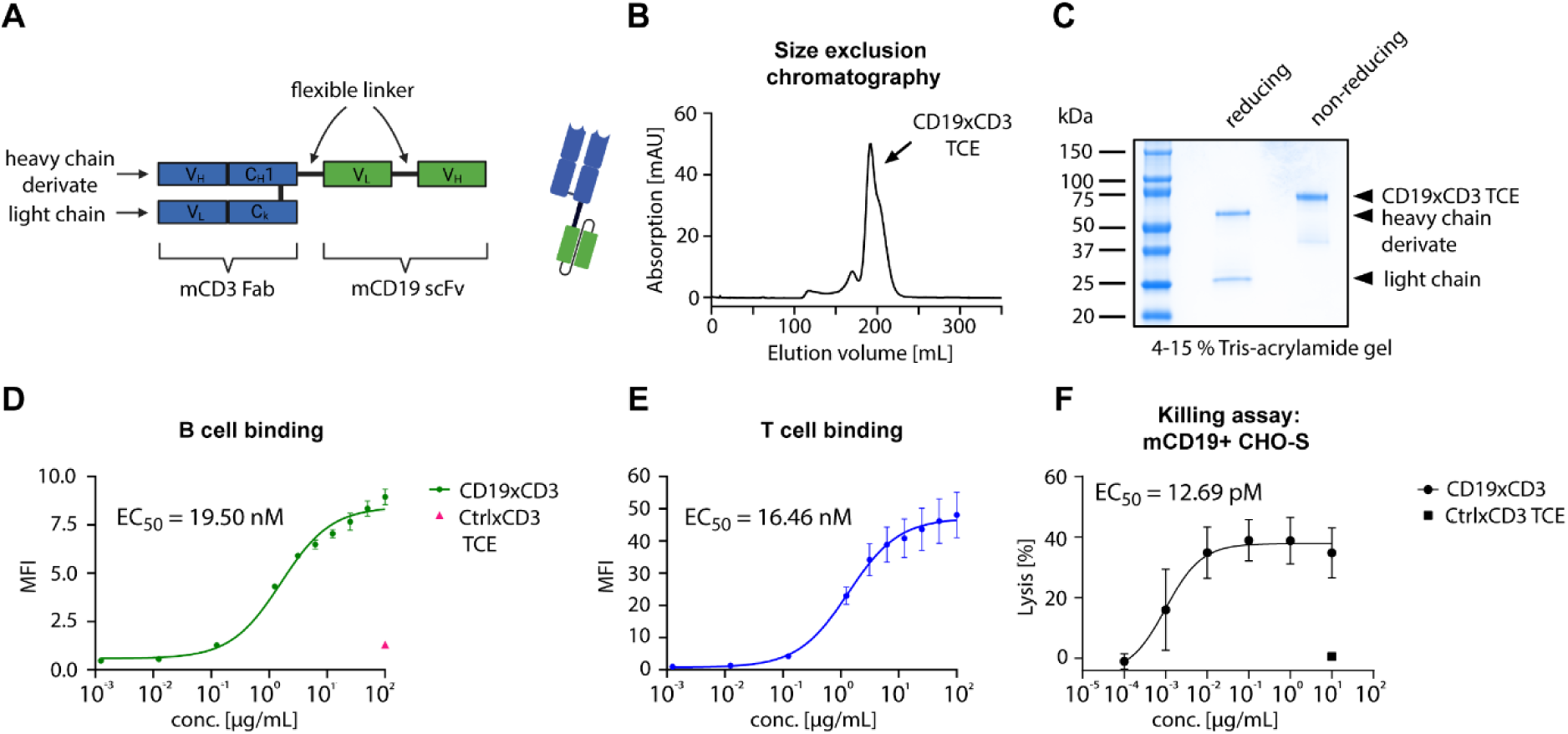
Design, production, and characterization of the CD19×CD3 TCE: The CD19×CD3 TCE was produced in CHO-S cells using the MaxCyte (STX) electroporation system. The molecule was purified from cell culture supernatants by affinity chromatography (CaptureSelect™ CH1-XL, ThermoFisher), followed by size exclusion chromatography (ÄKTA Pure). **(A)** Schematic representation of the CD19×CD3 TCE. **(B)** Size exclusion chromatography of the purified CD19×CD3 TCE. The peak at 190 mL represents the monomeric form of the molecule. **(C)** Coomassie blue-stained SDS-PAGE analysis under reducing and non-reducing conditions. **(D)** Concentration-dependent B cell binding, analyzed via flow cytometry. Murine B cells were isolated from the spleen using the Pan B Cell Isolation Kit (mouse, Miltenyi). **(E)** Concentration-dependent T cell binding, analyzed via flow cytometry. Murine T cells were isolated from the spleen using the Pan T Cell Isolation Kit II (mouse, Miltenyi). **(F)** CD19×CD3 TCE-mediated target cell killing, analyzed by a chromium release assay. Isolated T cells from murine spleens were used as effector cells at an effector-to-target ratio of 40:1. Transiently transfected CHO-S cells expressing murine CD19 were used as target cells. Data are presented as mean values ± SD (for binding curves) or ± SD (for killing assays) from three independent experiments. TCE: T cell enager.

### Doses of 1 µg CD19×CD3 TCE and above result in B cell depletion lasting 24 hours

Initial dose-finding was performed in healthy B6.s mice, with the endpoint being the relative number of B cells and T cells to the number prior injection, and the aim of reaching at least a 50 % reduction (**Figure 2A,B**). TCE were well tolerated, indicated by a low burden across all groups and no change in body weight (**Supplemental Table S1**). A 50 % reduction was achieved with a dose of 1 µg or higher irrespective of the application route (**Figure 2C**). Using the lowest concentration (0.1 µg), B cell counts decreased to 29 % (i.p) and 51 % (i.v). Higher doses than 10 µg only led to a slightly more pronounced depletion. Of note, frequencies of T cells (measured by CD4^+^ and CD8^+^ cells) were also decreased in peripheral blood with doses of 1 µg and above. With doses of 10 µg and higher, T cells remained depleted for 24 hours. Higher doses did not prolong the T cell depletion. However, irrespective of the dose and application route, the relative B cell count recovered after 2 days, indicating that multiple injections were needed for a sustained B cell depletion. Due to the greater burden of repetitive i.v. injections and similar depleting activates of the alternative application routes, i.p. injections were selected for the subsequent experiments.

**Figure 2:**
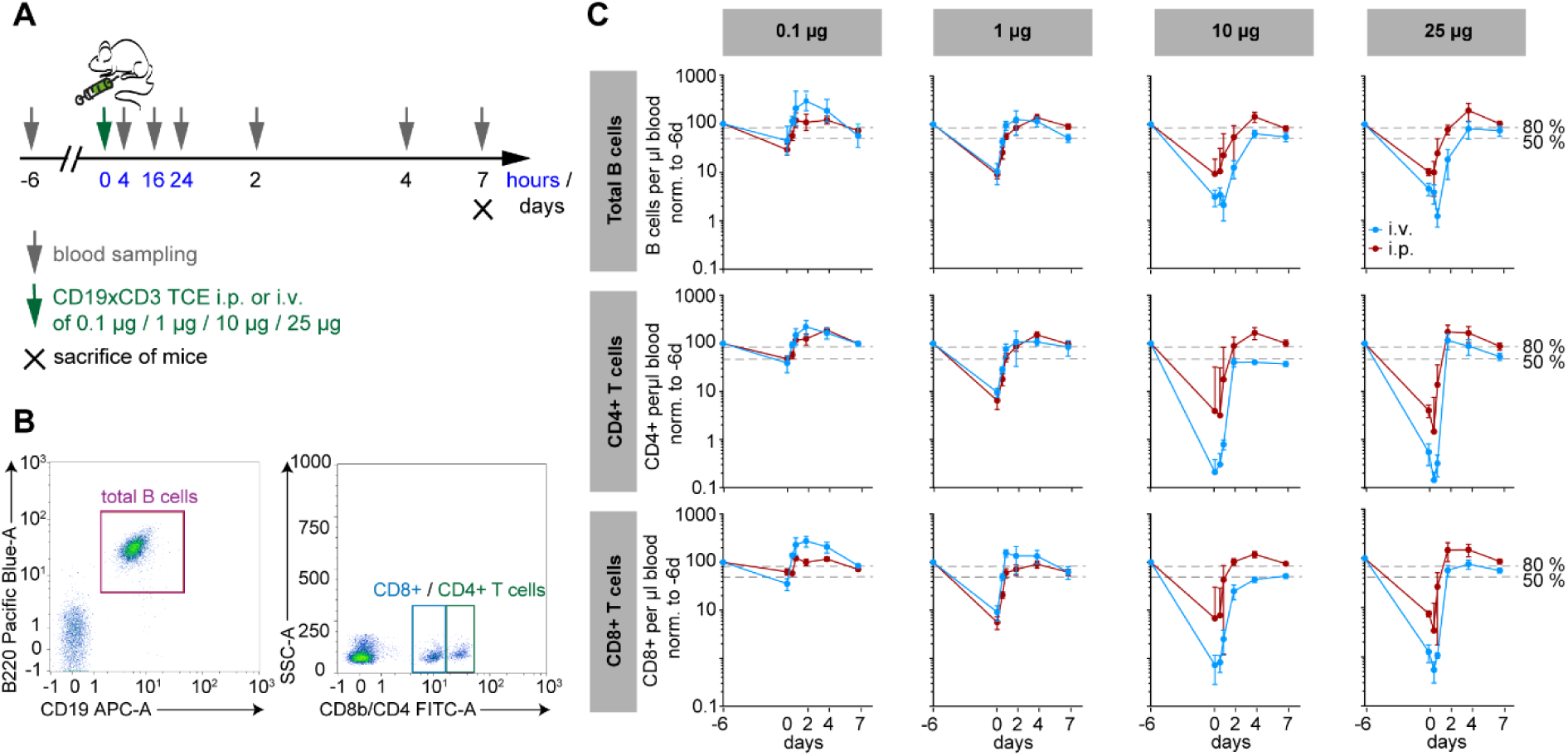
Efficient B cell depletion in peripheral blood with 1 µg of CD19×CD3 TCE. Dose finding was conducted in healthy B6.S mice to find the optimal dosage and route of application of the CD19×CD3 TCE to achieve a B cell depletion in peripheral blood, aiming a relative depletion of at least 50 %. **(A)** Schematic of the experimental setup with either intra-peritoneal (i.p.) or intra-venous (i.v.) injections in dosages of 0.1 µg, 1 µg, 10 µg, or 20 µg. Blood was taken from the submandibular vein 6 days before and 4, 16, and 24 hours, as well as 2, 4, and 7 days after the injection of the CD19×CD3 TCE. B and T cell frequencies were measured by flow cytometry. After one week, the experiment was terminated, and mice were euthanized. **(B)** Representative flow cytometry plots for total B cells (CD19^+^B220^+^) and T cells (CD8b^+^ or CD4^+^) from one mouse at time point 0. Both cell populations were gated from single living CD45^+^ cells and cytotoxic and helper T cells were distinguished from each other by the fluorescence intensity of the flow cytometry antibodies against CD8b and CD4. **(C)** Time course of the number of total B cells, CD4^+^, and CD8^+^ T cells per µl blood after the injection of the CD19×CD3 TCE either i.p. (red) or i.v. (blue) in four different dosages. Values were normalized for each mouse to the baseline level. Data are represented as mean values ± SD with n = 3 per group. TCE: T cell enager.

### Multiple injections of TCE lead to sustained B cell depletion in blood and bone marrow

To achieve a prolonged B cell depletion, healthy B6.s mice received additional i.p. injections of a dose of 1 µg CD19×CD3 TCE after 2 and 4 days (**Figure 3A**) and in comparison, to the previous experiment, spleen and bone marrow were additionally investigated. Weight loss and total burden were comparably low in all groups (**Supplemental Table S2)**. This time, a single injection of 1 µg TCE led to a more pronounced B cell depletion, lasting up to 4 days in the blood. The same was obtained for CD4 and CD8 T cells. Assessment of spleen and bone marrow showed no depletion of B or T cells using a single injection. In the peripheral blood, following the second injection on day 2, a more than 7-fold increase of B cells was observed. A third injection reduced B cell counts markedly to 24 % by day 7, remaining low thereafter. In the bone marrow, B cells decreased after the second injection and remained low throughout the experiment. These findings suggest a mobilization of B cells from the bone marrow into the circulation, potentially explaining the transient peripheral increase, where they were subsequently depleted by the TCE injection on day 4. (**Figure 3B**).

**Figure 3:**
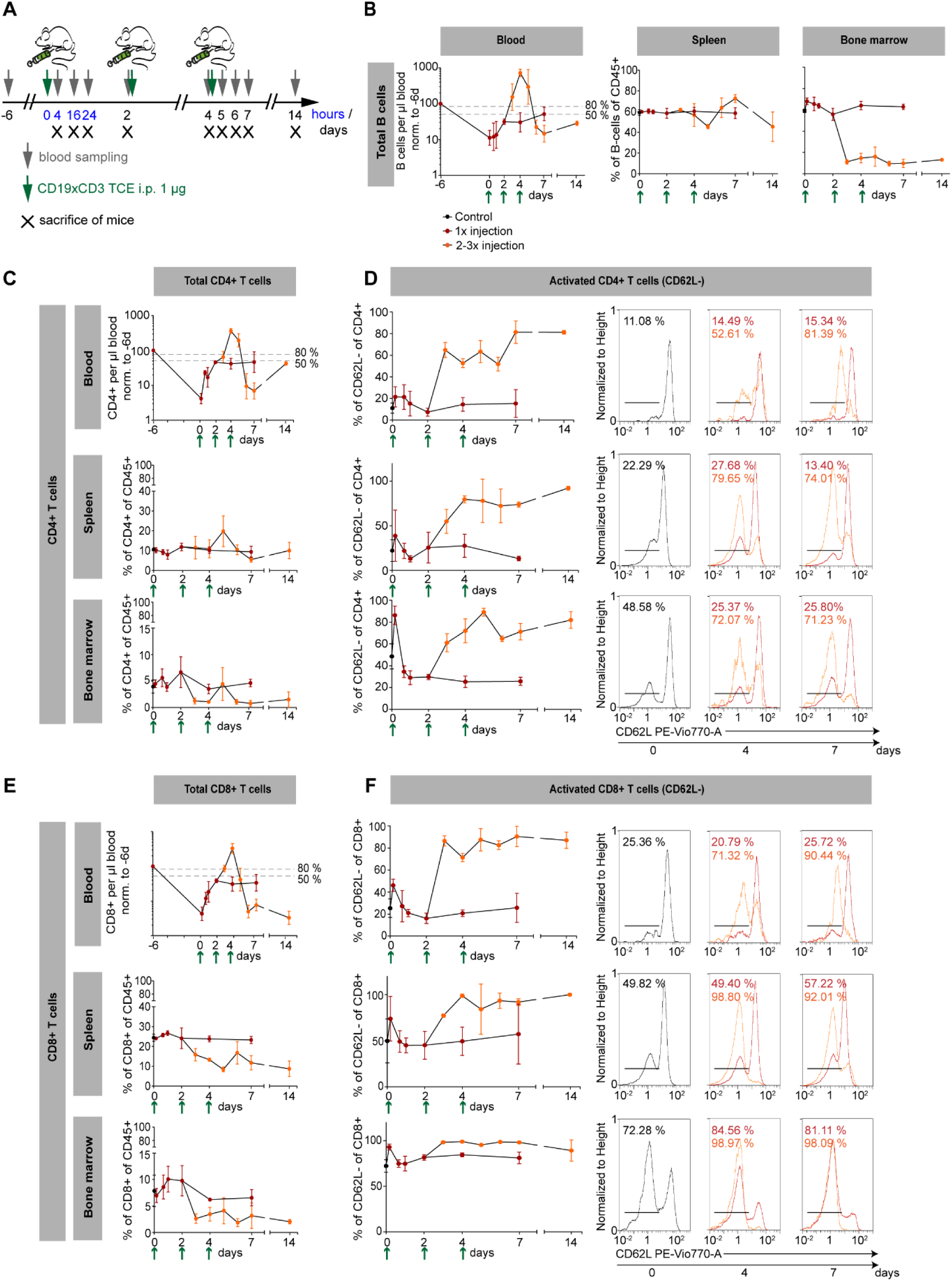
Consecutive injections of the CD19×CD3 TCE result in a stable depletion of B cells and activation of T cells in different compartments. For a deeper characterization of the effect of the CD19×CD3 TCE on B and T cells in blood, spleen, and bone marrow, the TCE construct was tested in healthy B6.S mice using repeated injections of 1 µg i.p.. **(A)** Schematic representation of the experiment: TCE were injected on day 0, 2, and 4 and blood sampling was performed 6 days prior, and 4, 16, 24 hours, and 2, 4, 7, and 14 days after the first injection. Mice were sacrificed at the indicated time points to analyze the B and T cells in blood, spleen and bone marrow. **(B, C, E)** Comparison of the effect of a 1x injection (red) to 2-3x injection (orange) at the given time points (green arrows) on **(B)** total B cells, **(C)** CD4^+^ and **(E)** CD8^+^ T cells in blood, spleen, and bone marrow. Values in blood were normalized to the baseline level for each mouse obtained from the first blood sampling 6 days prior of the first injection. In spleen and bone marrow, B and T cells are depicted as percentages of CD45^+^ cells. **(D, F)** For T cells, activation was assessed by the shedding of CD62L (CD62L^-^ cells) by flow cytometry for **(D)** T helper and **(F)** cytotoxic T cells in the respective lymphatic organs and compared between the 1x and 2-3x injection group. Respective histograms with marked CD62L^-^ population are shown for day 0 (control group), as well as day 4 and 7 with the mean percentage of CD62L^-^ cells for each group. Activated CD62L^-^ T cells are depicted as percentage of CD4^+^ or CD8^+^ T cells. Untreated healthy B6.S mice were used to obtain control values for spleen and bone marrow and data are represented as mean values ± SD with n = 3 per group. TCE: T cell enager.

Similar to the findings obtained for B cells, the second injection led to an increase in T cells in the peripheral blood, with a more than 3-fold increase of CD4 and CD8 T cells on day 4 (**Figure 3C**). In the bone marrow, CD4 T cells were persistently decreased from day 7 onward. CD8 T cells in spleen and bone marrow decreased following the second and third injections. These findings suggest that, as with B cells, T cells may transiently migrate from lymphoid tissues into the circulation following the second injection, where they are subsequently depleted by the repeated TCE administration (**Figure 3C, E**). Single injections of CD19×CD3 TCE led to a relatively short activation of CD4 T cells in all analyzed organs, measured by shedding of the CD62L antigen (**Figure 3D**). In contrast, after second and third injections, levels of activated CD4 T cells increased to up to 8-times higher levels relative to before TCE injection. Similar results were obtained for CD8 activated T cells in analyzed organs (**Figure 3F**).

In summary, these experiments demonstrate that, while B cell depletion in the peripheral blood was achieved using a single injection of 1 µg of TCE, additional depletion in the bone marrow could only be obtained through repeated injections after 2 and 4 days. Therefore, for the following experiments, 1 µg administered twice times weekly was selected as the appropriate dosage to achieve a sustained B cell depletion across compartments.

### Immunization-induced EBA is improved by therapeutic treatment with CD19×CD3 TCE

Next, we assessed the clinical efficacy of CD19×CD3 TCE in the immunization-induced EBA mouse model. Following EBA induction, mice reaching an affected body surface area (ABSA) of ≥ 2 % were randomized to receive either PBS or 1 µg of TCE twice weekly. Unexpectedly, no depletion of circulating B cells was noted. Hence, after 3-5 weeks of treatment, dosage was increased to 10 µg. Treatment was continued for up to 13 weeks or until ABSA dropped below 1 % for two consecutive weeks, indicating remission. At the end of the experiment, mice were euthanized and organs were taken for further analysis (**Figure 4A**).

**Figure 4:**
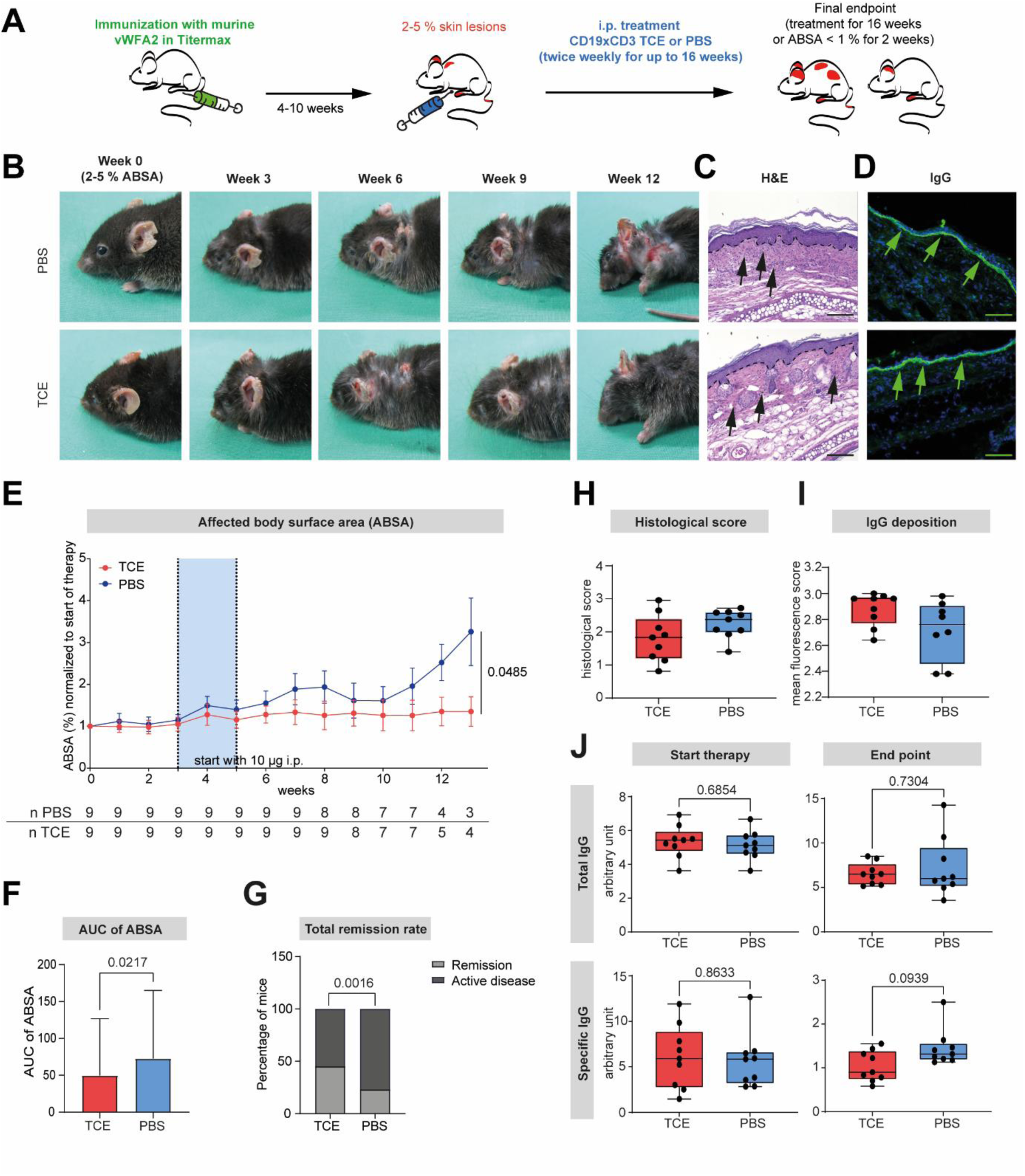
Treatment with CD19×CD3 TCE inhibits disease progression and promotes remission in an immunized mouse model of epidermolysis bullosa acquisita (EBA) To evaluate the clinical efficacy of the CD19×CD3 TCE, healthy B6.S mice were immunized on day 0 by injecting 60 µg murine vWFA2 antigen dissolved in Titermax® in both hind foot pads **(A)**. EBA skin lesions developed after 3-5 weeks and mice were weekly scored to assess the percentage of the affected body surface area (ABSA). If ABSA increase above 2 %, mice were randomized to the treatment (CD19×CD3 TCE, i.p.) or control group (PBS, i.p.). Treatment was performed twice weekly for up to 13 weeks, or until mice reached remission (ABSA < 1 % for two consecutive weeks). **(B)** Representative pictures of both groups at day of randomization, and after 3, 6, 9, and 12 weeks of treatment. **(C)** Representative pictures of a H&E-stained histological section from the ear of one mouse of each group at the end of the experiment. The dotted line indicated the dermal-epidermal junction (DEJ) and black arrows show infiltrating immune cells in the dermis. **(D)** Direct immunofluorescence of IgG deposition (green arrows) at the DEJ on ear sections. **(E)** ABSA) over time, normalized for each mouse to the randomization day, is shown with the treatment group (TCE) in red and the control group (PBS) in blue. Treatment started with 1 µg of TCE twice a week and between week 3 and 5, dosage of TCE was increased to 10 µg per injection and continued until the end of the experiment. Below, number of mice per group at each timepoint indicated. **(F)** Comparison of the area under the curve (AUC) of the ABSA from start of experiment until week 13. Analysis of AUC showed that mice treated with TCE exhibited a significant inhibition of disease progression. **(G)** Percentage of mice in remission of each group (TCE 45 %, PBS 23 %). **(H)** Histological score, including epidermal thickness, dermal infiltration and split formation, and **(I)** IgG deposition showed no difference between both groups. **(J)** Total and vWFA2 specific IgG levels in plasma at the start and end of therapy showed no significant difference. Data are presented as **(E, F)** mean values ± SD, **(H-J)** medians (black line), 25^th^/75^th^ percentiles (boxes), and max/min values (error bars) or **(G)** stacked bars with n = 3-9 per group. Statistical analysis: **(E)** Two-way ANOVA with mixed-effects model, **(F)** unpaired t-test, **(H-J)** Mann-Whitney test, and **(G)** Fisher’s exact test. Scale bar: 100 µm. TCE: T cell enager.

In line with previous experiments, the first signs of disease were observed three weeks post-immunization.^32–34^ In total, 18 mice met the inclusion criteria, leading to randomization of 9 mice per group. There was no difference in change of body weight between the two groups throughout the whole experiment. Total burden increased irrespective of the treatment **(Supplemental Figure S2)**. Mice receiving PBS showed a continuous increase in ABSA (normalized to the day of randomization), reaching a maximum mean of 3.26 % after 13 weeks. In contrast, mice treated with TCE exhibited only a marginal increase, with a peak mean ABSA of 1.35 %. Overall, analysis of area under the curve (AUC) showed a significant difference of the ABSA between the two groups (**Figure 4B, E, F**). Twice as many mice achieved disease remission in the TCE group (4/9; 45 %) compared to the PBS group (2/9; 23 %) (**Figure 4G**). Early euthanasia due to disease burden was required in 3 of 9 mice (34 %) receiving TCE, compared to 5 of 9 mice (56%) in the PBS group. One mouse in each group died under anesthesia, and one mouse per group completed the full 16-week treatment period (**Table 1**). In summary, treatment with CD19×CD3 TCE resulted in improved clinical outcomes, including higher remission rates, reduced disease progression, and decreased need for early euthanasia, supporting their therapeutic potential in pemphigoid diseases.

**Table 1:**
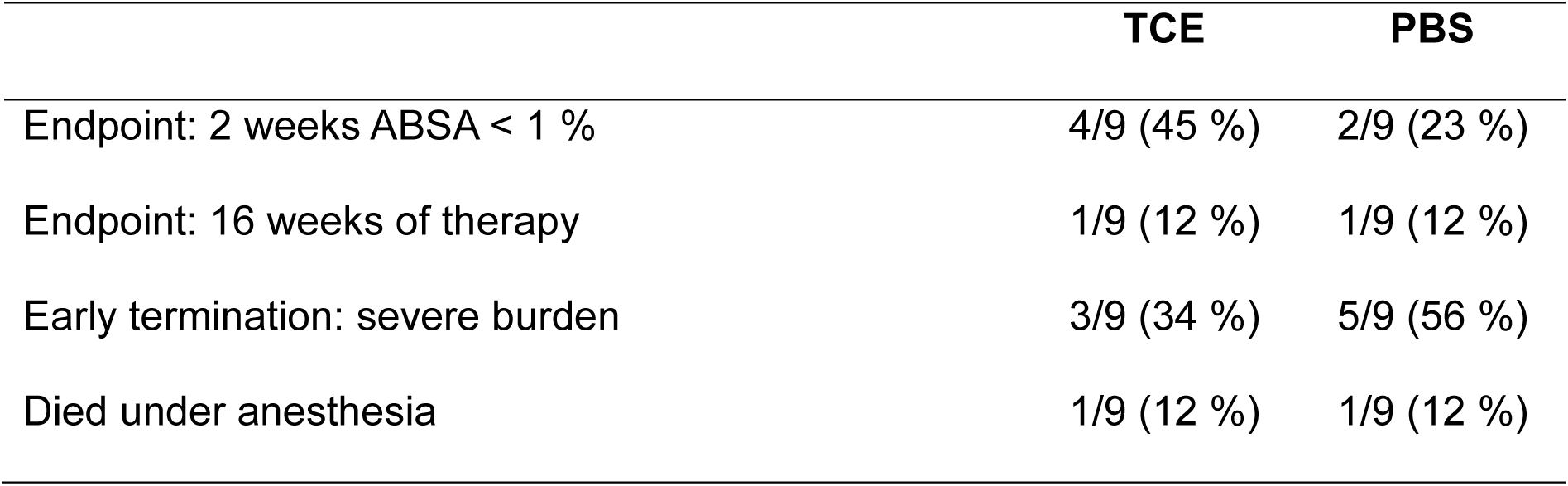
Number of mice reaching final endpoints or undergoing early termination. Two endpoints were defined after randomization: mice either went into remission, defined as an ABSA score below 1 % for two consecutive weeks, or completion of the 16-week treatment period. Mice were subject to early termination if they exhibited a high total burden (weight loss, change in behavior and physical appearance, and ABSA score). In both groups, 9 mice were successfully randomized. The data include both the absolute number and the relative percentage of mice. TCE: T cell enager.

### Comparable histological findings in H&E staining and IgG deposition

At the indicated end point, skin samples were taken for further analysis. Sections where stained by H&E and infiltration and split formation was assessed, but did not differ between groups (**Figure 4C and H**). Direct immunofluorescence staining for IgG was performed, the mean fluorescent score did not differ between both groups (**Figure 4D and I**). Overall, histological analysis of H&E staining and IgG and C3 (**Supplemental Figure S**3) deposition revealed similar findings in both groups.

### TCE-treatment does not affect specific vWFA and total IgG levels

Total IgG and anti-mCOL7^vWFA2^ IgG levels were measured in plasma samples throughout the treatment period. At randomization, total IgG levels were comparable between groups (5.34 U (0.95) in TCE-treated mice and 5.13 U (0.86) in PBS controls). By the individual endpoint of each mouse, total IgG levels had increased slightly in both groups, reaching 6.53 U (1.21) (TCE) and 7.22 U (3.3) (PBS). Anti-mCOL7^vWFA2^ IgG levels on day 0 were 6.06 U (3.46) in the TCE group and 5.64 U (3.58) in PBS controls. These levels declined in both groups by the study endpoint, to 1.04 U (0.35) and 1.46 U (0.42), respectively (**Figure 4J**). Taken together, CD19×CD3 TCE treatment was associated with a modest, not statistically significant, reduction in pathogenic Aab, while total IgG levels remained unaffected.

### Specific vWFA2 B cells are depleted by TCE-treatment in the blood

To assess the effects of CD19×CD3 TCE treatment on B and T cell frequencies, we performed flow cytometric analysis of blood, spleen, and bone marrow. Notably, treatment reduced circulating vWFA2-specific B cells (**Figure 5A-B**). At the endpoint, relative vWFA2-specific B cell counts were 56.3 % (83.9) in TCE-treated mice versus 258.0 % (231.0) in PBS controls, compared to day 0. Total and antigen-specific B cell counts in other compartments remained unchanged. Among T cells, CD4⁺ T cell frequencies were reduced in mesenteric lymph nodes (9.6 % (5.2) in TCE-treated vs. 18.3 % (9.5) in PBS) and spleen (4.2 % (1.5) vs. 7.2 % (3.1), respectively). No significant differences were observed in other compartments (**Figure 5C, D**). Overall, CD19×CD3 TCE treatment selectively depleted antigen-specific B cells in peripheral blood and reduced CD4⁺ T cell frequencies in secondary lymphoid tissues.

**Figure 5:**
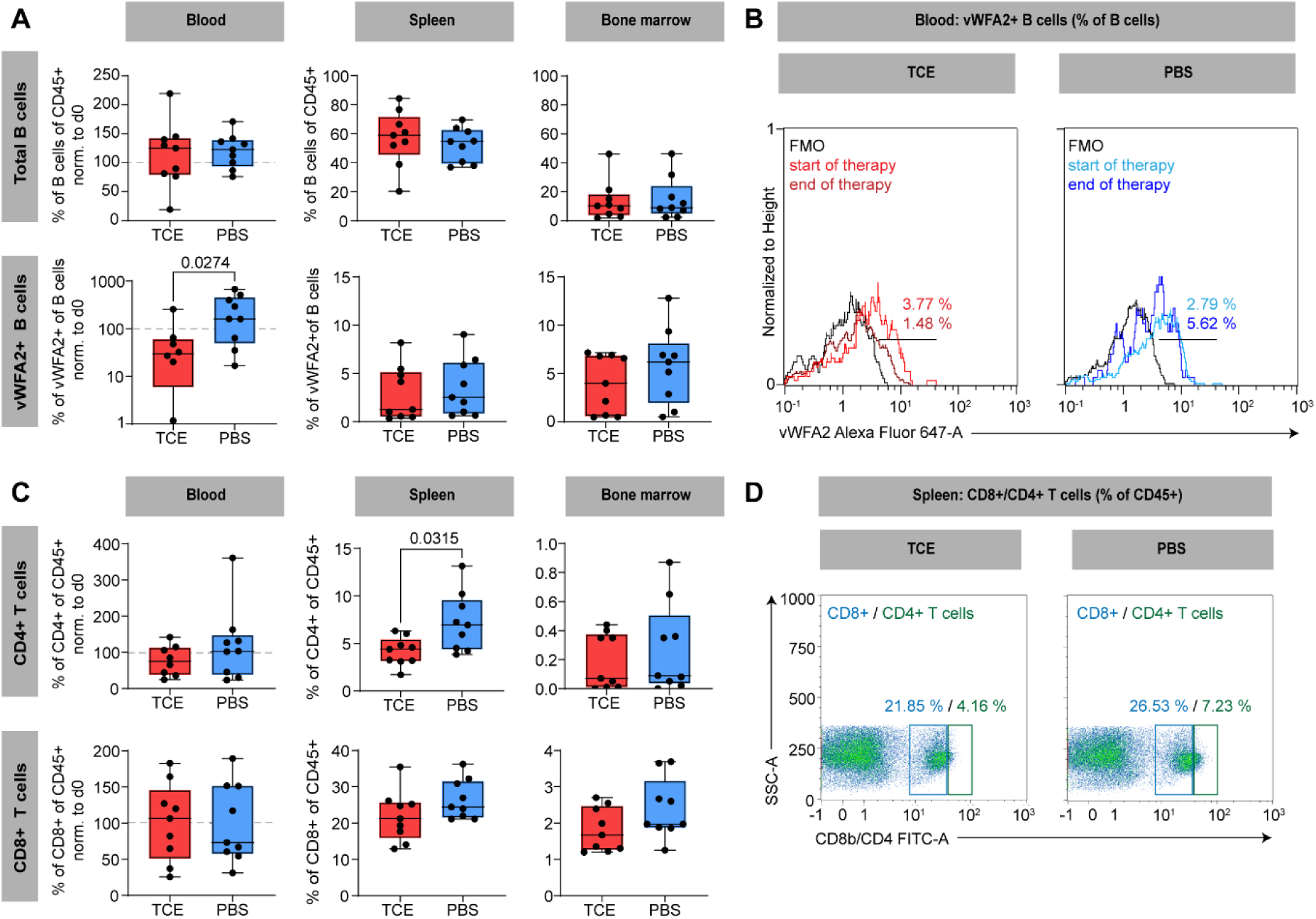
Levels of specific B cells are significantly reduced in peripheral blood after treatment with CD19×CD3 TCE. Total and vWFA2-specific (vWFA2^+^) B cells, as well as T helper and cytotoxic T cells from various lymphatic organs were analyzed by flow cytometry in mice treated with TCE in the immunization-induced model of EBA. **(A)** Percentages of total or vWFA2^+^ cells in blood, spleen, and bone marrow of both respective groups. Total B cells are defined as CD19^+^B220^+^ and are depicted as percentages of CD45^+^ cells and vWFA2^+^ B cells are shown as percentages of total B cells. Values for blood are normalized to the individual values on day of randomization. In blood, TCE treatment lead to significant fewer vWFA2^+^ B cells than the control group. **(B)** Flow cytometric histograms of vWFA2^+^ B cells in blood normalized to height. For each group, the plot shows the vWFA2^+^ B cell population at start of therapy (randomization day) and end of therapy as well as the corresponding FMO control. Depicted are the mean values of the vWFA2^+^ B cells. **(C)** Percentages of T helper (CD4^+^) or cytotoxic T (CD8^+^) cells in blood, spleen, and bone marrow of both respective groups. Both T cell populations are shown as percentages of CD45^+^ cells and values for blood are normalized to day of randomization. In TCE-treated mice, CD4^+^ T cells were significantly reduced in spleen. **(D)** Representative flow cytometric density plots of CD8^+^ and CD4^+^ T cells gated from CD45^+^ cells of both groups. Shown are the mean percentages of T helper and cytotoxic T cells. Data are presented as medians (black line), 25^th^/75^th^ percentiles (boxes), and max/min values (error bars) with n = 7-9 per group. Statistical analysis: Mann-Whitney test. TCE: T cell enager.

## Discussion

Our study demonstrates that a CD19×CD3 TCE at doses of 1 µg and above effectively depletes B cells in mice under steady-state conditions, with effects lasting several days. Repeated administration led to sustained B cell depletion in both blood and bone marrow, indicating potential for long-term, compartment-selective B cell depletion. Given the central role of autoreactive B cells in PD, we wanted to investigate whether the bispecific T cell engager-mediated depletion translates into therapeutic benefit in a disease model. In the immunization-induced EBA model, therapeutic TCE treatment significantly improved clinical outcomes, highlighting its efficacy in autoimmune diseases. Notably, TCE treatment also depleted circulating vWFA2-specific B cells. Taken together, these findings support the potential of CD19×CD3 TCE as a therapeutic strategy for AAb-mediated diseases, under the conditions modeled in the induced murine EBA.

In the initial studies that led to the FDA approval of blinatumomab for Philadelphia chromosome–negative relapsed or refractory precursor B-cell acute lymphoblastic leukemia (R/R B-ALL), blinatumomab was continuously given as an intravenous infusion over 4 weeks of a 6-week cycle.^33,34^ A step-dose increase from 5 µg to 15 μg/m^2^/day was found to be best tolerated.^34^ Blinatumomab and the CD19×CD3 TCE used in this study are both designed without an Fc part, which leads to significantly restricted pharmacological properties. Due to their low molecular mass (55 kDa for blinatumomab, 77.6 kDa for the CD19×CD3 TCE) and the absence of FcRn-mediated recycling, both antibodies exhibit a serum half-life of only a few hours^35^. As a consequence, blinatumomab is clinically administered via continuous infusion to reduce fluctuating systemic concentrations, avoiding the risks associated with high-dose bolus injections, such as CRS or ICANS^36^. However, hematological malignancies such as B-ALL typically involve high burdens of tumor/target cells, necessitating sustained levels of antibody for high therapeutic efficacy. In contrast, autoreactive B cells in autoimmune diseases such as EBA represent only a minor fraction of the total B cell pool, which may allow for effective depletion without continuous infusion. In our initial dose-finding experiments that administered the TCE via intraperitoneal injections three times per week, we demonstrated that this treatment regimen is sufficient to achieve B cell depletion not only in blood but also in the bone marrow. Additionally, dosages of up to 25 µg per infusion were well tolerated and did not lead to weight loss or signs of toxicity in mice. Unlike in hematological cancer treatment, where continuous antibody presence is essential, the intermittent administration of CD19×CD3 TCE appears sufficient for effective modulation of (autoreactive) B cells in autoimmunity settings.

Treatment with CD19×CD3 TCE alleviated immunization-induced EBA. Unexpectedly, total B cell counts remained largely unchanged despite clinical improvement. Notably, there was a reduction in vWFA2-specific B cells. This might be related to differences in CD19 expression levels or a higher susceptibility to T cell-mediated killing. The lack of sustained B cell depletion may be attributable to the use of TiterMax^®^, an adjuvant essential for disease induction in this model. TiterMax^®^ contains metabolizable oil, block copolymer CRL-8941, squalene, and a microparticulate stabilizer^36^ and induces not only the generation of mCOL7-specific B cells but also a robust local immune response. Pronounced swelling of inguinal lymph nodes and footpads (unpublished) suggests heightened B cell activation and proliferation. Such an environment may exceed the depletion capacity of TCE, unlike in healthy mice, where a single injection is sufficient to markedly reduce B cell numbers. These findings suggest that in activated immune environments, such as those induced by TiterMax^®^ administration or during active disease flares, the depletion capacity of bispecific antibodies can be limited. This highlights the challenge of B cell-directed therapies in the context of ongoing immune activation and underscores the importance of evaluating treatment efficacy in disease-relevant models.

The limited pharmacological properties of Fc-lacking bispecific antibodies, especially the significantly reduced serum half-life, necessitate continuous administration for maintained therapeutic levels. To address this, most bispecific antibodies in clinical development are engineered as IgG-like molecules, like the BCMA×CD3 antibodies linvoseltamab and teclistamab, which benefit from prolonged bioavailability by FcRn-mediated recycling.^37,38^ However, beyond systemic persistence, other factors such as the ability of an antibody to penetrate tissues and reach target cells in their specific niches might be equally if not more critical, particularly in autoimmune diseases, where autoreactive B cells may reside in peripheral tissues such as lymphoid organs. In this context, smaller molecules such as the CD19×CD3 TCE may offer a distinct advantage in accessing otherwise poorly perfused tissues or compartments.^39^ Consequently, future therapeutic strategies should not only focus on clinical efficacy but also on understanding the biological underpinnings of each disease, including the localization and phenotype of pathogenic B cell populations. Comparative studies of both antibody formats (IgG-like vs. non-IgG-like) and B cell-targeting specificities will be crucial. Importantly, multiple bispecific T cell engagers that have received FDA approval for hematological malignancies are currently clinically repurposed for the treatment of autoimmune entities, such as teclistamab, highlighting the expanding therapeutic potential of this class of immunotherapies.^21^

Despite the observed clinical improvement in EBA mice, mCOL7^vWFA2^ Aab titers were unexpectedly not significantly reduced following CD19×CD3 TCE treatment. This finding is in line with the known resistance of long-lived plasma cells to CD19-targeted therapies. As terminally differentiated B cells, plasma cells typically lack surface expression of CD19 and are thus not targeted or eliminated by CD19×CD3 bispecific T cell engagers.^40–42^ In addition, previous studies have shown that plasma cells induced by vaccination can persist for over a decade, and CD20-directed therapies do not affect pre-existing humoral immunity, so that antibody titers against tetanus, measles, mumps, and rubella remain unchanged.^43^ Although CD19 is expressed on a subset of plasmablasts and early plasma cells^44^, targeting CD19 alone may not suffice to eliminate the full pool of Aab-producing cells. These findings suggest that combining CD19-directed therapies with other molecules targeting plasma cell antigens, such as BCMA, could further enhance therapeutic efficacy in Aab driven diseases. Recent clinical case reports also support this combinatorial approach^45^, a strategy that can be tested in subsequent studies.

In summary, we demonstrate that selective B cell depletion using CD19×CD3 TCE is a promising strategy for treating Aab-mediated autoimmune diseases such as PD. Moreover, the therapy was well tolerated and could potentially be applied even in vulnerable patients. Our study may serve as a blueprint for related conditions, as the antigen specificity of TCE can, in principle, be adapted to match the unique immunopathology of PD and other autoimmune diseases.

## Materials and Methods

### Design, expression and purification of mCD19×CD3 TCE

For the generation of the bispecific CD19×CD3 TCE in a bibody format, two plasmids for secretion (pSEC) encoding the heavy chain derivative and the light chain were designed. For the heavy chain derivative, the variable heavy chain (VH) of the anti-mCD3 antibody clone 145-2C11^46^ and the constant heavy 1 (CH1) domain were fused to an anti-mCD19 scFv coding sequence (clone 1D3) via two flexible glycine-serine linkers. Plasmid DNA was amplified and purified using the Nucleo Bond 2000 EF (Macherey-Nagel, Dürren, Deutschland) and the correct sequences were confirmed by Sanger sequencing (Institute of Clinical Molecular Biology, Kiel, Germany). The TCE was produced in CHO-S cells using the MaxCyte STX large scale electroporation system, following the manufacturer’s recommendations. For purification from cell culture supernatant, the Capture Select IgG-CH1-XL affinity matrix (ThermoFisher) was used, followed by size exclusion chromatography performed with the ÄKTA pure system (Cytiva). As an isotype control, an antibody targeting murine CD3 (clone 145-2C11) and an irrelevant antigen (Ctrl**×**CD3) was generated in the same molecular format.

### SDS-PAGE analysis

Two micrograms (µg) of purified protein were loaded on a precast 4-15% Tris-acrylamide gel (Mini-PROTEAN^®^ TXN^™^, BioRad) under reducing and non-reducing conditions and stained overnight with Coomassie Brilliant Blue staining solution (Carl Roth GmbH).

### Transient transfection of CHO-S cells

For *in vitro* cytotoxicity assays, murine CD19-expressing CHO-S cells were generated via transient transfection. Therefore, the MaxCyte STX large scale electroporation system was used following the manufacturer’s recommendations. 24 h post-electroporation, surface expression of murine CD19 was verified by flow cytometry.

### Assessment of in vitro cytotoxicity

Cytotoxicity assays were performed as described.^47^ In brief, target cells were labeled with 50 µCi of ^51^Cr for 2 h followed by three washes with culture medium. For effector cell preparation, murine T cells were isolated from spleen using the Pan T cell Isolation kit (Mouse) (Miltenyi Biotec) following the manufacturer’s recommendations. Effector cells, culture medium and antibody were added to a round bottom microtiter plate. The assays were initiated by dispensing the labeled target cells, i.e. murine CD19-expressing CHO-S cells, at an effector to target (E:T) cell ratio of 40:1 in a total volume of 200 µL. After incubation for 18 h at 37°C, ^51^Cr release from triplicates samples was measured in counts per minute (cpm) using the MicroBeta TriLux 1450 liquid scintillation and luminescence counter (PerkinElmer/Revvity). Percentage of cellular cytotoxicity was calculated using the formula: % specific lysis = (experimental cpm – basal cpm) / (maximal cpm – basal cpm) x 100. Maximal cpm was determined by adding Triton X-100 to target cells and basal release was determined in the absence of antibody or effector cells.

### Animal experiments

Immunization-EBA susceptible B6.SJL-H2s C3c/1CyJ (B6.S) mice ^31^ were originally obtained from the Jackson Laboratories (Stock 000966; Bar Harbor, Maine, USA) and bred at the animal facility of the University of Lübeck, Germany. For the experiments, sex-matched adult mice (>8 weeks old) were used. The mice were given standardized mouse chow as well as acidified drinking water provided ad libitum and maintained on a 12-hour light/dark cycle. All clinical examinations were performed under anesthesia, using intraperitoneal (i.p.) administration of a ketamine (100 µg/g body weight, Sigma-Aldrich, Germany) and xylazine (15 µg/g body weight, Sigma-Aldrich, Germany) mixture unless otherwise specified. Blood sampling was performed on awake animals by puncture of the vena facialis following the recommendations of the GV-SOLAS (Freiburg, Germany). Mice were euthanized under deep anesthesia, followed by complete exsanguination and cervical dislocation. All animal experiments were conducted as per the European Community rules for animal care and were approved by the governmental administration (V242-49267/2019 (80-7/19), Ministry for Energy, Agriculture, the Environmental and Rural Areas) and conducted by certified personnel.

### Dose-finding studies of TCE in healthy mice

Dosing was performed in healthy B6.S mice with n=3 per group. After initial blood sampling 6 days prior, CD19×CD3 TCE were dissolved in phosphate-buffered saline (PBS) and injected in four different concentrations (0.1, 1, 10 and 25 µg per mouse) either i.p. or i.v.. Weight was recorded at day −6, 0, and at the end of the experiment with day 0 being the day of TCE administration. Blood was also taken at day −6 and after 4, 16, 24 hours and 2, 4 and 7 days after antibody injection. After 7 days mice were euthanized. In a second set of experiments, blood sampling was again performed at the previously indicated time points, this time extended to 14 days. Application of 1 µg of TCE was repeated after 48 and 96 hours. In this experiment mice were euthanized at different time points to investigate B cell depletion across different immunological compartments.

### Immunization-induced EBA and pre-clinical study protocol

For induction of experimental EBA, B6.S mice were immunized with 120 μg of the vWFA2 domain of COL7 (mCOL7^vWFA2^) mixed in a 1:1 ratio with TiterMax^®^ by injecting 60 μL of the vWFA2/TiterMax^®^ mixture into each footpad of the hind legs, as previously published ^48^. Clinical evaluations and total burden score (considering weight loss, changes in behavior and physical appearance), and clinical EBA severity, expressed as affected body surface area (ABSA), were carried out weekly by a person unaware of the treatment.^49^ When, within 10 weeks following immunization, mice reached ABSA-score of ≥ 2 %, they were randomly assigned to either the treatment group (receiving TCE; n=9) or the control group (receiving PBS; n=9). The TCE group was treated with i.p. injections of 1 µg per injection, and within 3-5 weeks dose was increased to 10 µg. Treatment was performed twice weekly for up to 16 weeks or until mice reached remission, classified as an ABSA < 1 % for two consecutive weeks. Blood samples were taken one week prior to immunization, at randomization, and every bi-weekly after randomization. After 13 weeks, all remaining mice (n=5) were euthanized and blood, skin and organs were taken for further evaluation. For assessment, relative clinical scores were calculated by comparing each week’s score to the initial score at the start of treatment (weekly score/initial score), overall disease severity was quantified as the cumulative relative clinical score over time (area under the curve), and remission rates were determined.

### Histology

For histology, skin and ear samples were fixed in 3.7 % paraformaldehyde and embedded in paraffin. Hematoxylin and eosin (H&E) staining was performed on 4-5 µm thick sections following standard protocols. The histological score was assessed by combining the individual epidermal thickness as well as split formation and infiltration of immune cells by an investigator unaware of the applied treatments.^50^ To detect tissue bound rabbit IgG and murine C3, direct immunofluorescence microscopy was performed as described.^6^ Briefly, frozen sections were prepared from 6 µm thick skin and ear sections and incubated with goat anti-rabbit antibodies reactive with rabbit IgG (Dako Deutschland GmbH, Hamburg, Germany) and murine C3 (MP Biomedicals LLC, Keyseberg, France). Sections were labeled with fluorescein isothiocyanate (FITC) and the respective fluorescence intensity was evaluated by a blinded investigator using a semi-quantitative assessment method.

### Flow Cytometry

Flow cytometry analysis was performed in independent experiments using either a Navios flow cytometer (Beckman Coulter) with data analyzed in Kaluza Analysis software (Beckman Coulter), or a MACSQuant Analyzer 10 (Miltenyi Biotec; Bergisch Gladbach, Germany) with data analyzed in MACSQuantify Software (Version 2.13; Miltenyi Biotec). Each experiment was conducted entirely on one of the two platforms. For concentration-dependent binding analysis of the CD19×CD3 TCE, 2 x 10^5^ MACS isolated murine B or T cells (C57BL/6) were washed in PBS containing 1% bovine serum albumin (BSA) an 0.1 % sodium azide (PBA buffer). Next, cells were incubated with the protein at varying concentrations on ice for 60 min and washed three times with 200 µL PBA buffer. Finally, cells were stained with a secondary anti-human-kappa-FITC antibody (SouthernBiotech) on ice for 30 min, washed three times with 200 µl PBA buffer and resuspended in 300 µL PBA buffer. For flow cytometric analyses of cell composition of blood, spleen, and bone marrow, single cell suspensions were prepared: in short, spleens were gently grinded between two slides in DPBS (Life technologies, Carlsbad, CA, USA) and cells were filtered through a 70 µm cell strainer. Bone marrow was isolated from femur and tibia, and cells were flushed out using a 27 G syringe and PBA buffer.^51^ Erythrocytes were lysed and life/dead staining was performed using the fixable viability dyes BD620 or 780 (BD Bioscience, Becton, New Jersey, USA). After a Fc-blocking step (Miltenyi Biotec) of 10 minutes on ice, cells were stained for 20 minutes with the following antibodies: Pacific-Blue, VioGreen, FITC, PE-Vio770, and APC-conjugated anti-mouse B220 (clone: RA3-6B2), CD45 (30F11), CD4 (GK1.5), CD8 (H35-17.2), CD62L (MEL14-H2.100), and CD19 (REA749), from Biolegend (San Diego, California, USA) or Miltenyi Biotec. Cells were gated for leukocytes (FSC-A and SSC-A) and singlets (FSC-H compared with FSC-A). Single cells were then differentiated between alive and dead. Only living cells were further gated on their positive CD45 expression and then the positive B cell (B220 and CD19) and T cell (CD4 and CD8) population, as well as activated T cells (CD62L negative) (Supplemental Figure S1). For specific vWFA2^+^ B cells, recombinant mCOL7^vWFA2^ was labeled with an Antibody Labeling Kit Alexa Fluor 647 (Invitrogen, Waltham, MA, USA) following manufacturès instruction. For flow cytometric staining, labeled mCOL7^vWFA2^-Alexa Fluor 647 was used in a pre-tested concentration alongside the above listed antibodies and specific B cells were gated from the positive B cell population. Positive vWFA2^+^ B cells were determined by comparing the fluorescence signal to a FMO control from the corresponding mouse.

### ELISA

Plasma levels of circulating total and autoreactive anti-mCOL7^vWFA2^ IgG of immunized mice were determined by ELISA using mouse quantification sets (Bethyl, Montgomery, Texas, USA) following the manufacture’s protocol with minor modifications for the autoreactive antibodies.^52,53^ In detail, each well was coated with 250 ng recombinant mCOL7^vWFA2^ in coating buffer. After blocking, diluted samples were added and incubated for 60 min. Bound antibodies were detected by HRP-conjugated goat anti-mouse antibodies (Bethyl) and tetramethylbenzidine (InvitrogenWaltham, MA, USA). The enzymatic color reaction was stopped by adding 2 M sulfuric acid (Carl Roth, Karlsruhe, Germany), and the change in OD was measured with a GloMax® Discover Microplate Reader photometer (Promega, Walldorf, Germany) at 450 nm. Standard reference curves were established by using the provided mouse reference sera (Bethyl).

### Statistical analysis

For graphical and statistical analyses, GraphPad Prism 10 software (GraphPad Software Inc., San Diego, CA, USA) was used. Applied tests and confidence intervals are indicated in the respective text and figure legend. A p-value < 0.05 was considered statistically significant. Sample size calculation for animal experiment were performed with SigmaPlot 13.0 (Systat Software Inc.) considering the percentage of with EBA affected body area at the end of the experiment as primary endpoint as well as the remission rates. The calculations assumed an expected standard deviation of 45 % in the immunization-induced EBA model, an alpha level of 5 %, a power of 85 %, and a minimum detectable difference from the positive control of 50 %.

## Data availability statement

The data that support the findings of this study are available from the corresponding author upon reasonable request.

## Acknowledgements

We thank the Institute of Clinical Molecular Biology in Kiel for providing Sanger sequencing as supported in part by the DFG Clusters of Excellence “Precision Medicine in Chronic Inflammation” and “ROOTS”. We thank T. Naujoks, Dr. D. Langfeldt and Dr. B. Löscher for technical support. The graphical abstract was created using BioRender.com.

## CRediT author statement

Natalie Gross - Investigation, Formal analysis, Visualization, Writing (initial draft)

Alexander Jochimsen - Investigation, Formal analysis, Writing (initial draft)

Sören Dräger - Investigation, Formal analysis, Writing (initial draft)

Thomas Theocharis – Investigtation, Writing (critical revision)

Andreas Recke – Resources, Writing (critical revision)

Nina van Beek – Writing (critical revision)

Leon Schmidt-Jiménez^1^ - Writing (critical revision)

Steffen Krohn – Writing (critical revision)

Katja Klausz – Methodology, Supervision, Writing (critical revision)

Katja Bieber – Conceptualization, Investigation, Methodology, Supervision, Writing (critical revision)

Matthias Peipp – Conceptualization, Resources, Methodology, Funding acquisition, Supervision, Writing (critical revision)

Ralf J. Ludwig - Conceptualization, Resources, Methodology, Funding acquisition, Supervision, Writing (critical revision)

## Declaration of generative AI and AI-assisted technologies in the writing process

During the preparation of this work the authors used ChatGPT 4o (OpenAI, San Francisco, CA, USA) in order to improve language and readability. After using this tool, the authors reviewed and edited the content as needed and take full responsibility for the content of the publication.

## Supplemental Information

Document Supplement: Tables S1 and S2, Figures S1-S3

## References

1. Hübner F, Kasperkiewicz M, Knuth-Rehr D, Shimanovich I, Hübner J, Süfke S, et al. Adjuvant treatment of severe/refractory bullous pemphigoid with protein A immunoadsorption. JDDG J Dtsch Dermatol Ges. 2018 Sept;16(9):1109–18.

2. Borradori L, Van Beek N, Feliciani C, Tedbirt B, Antiga E, Bergman R, et al. Updated S2k guidelines for the management of bullous pemphigoid initiated by the European Academy of Dermatology and Venereology (EADV). J Eur Acad Dermatol Venereol. 2022 Oct;36(10):1689–704.

3. Hoffmann JHO, Enk AH. High-Dose Intravenous Immunoglobulin in Skin Autoimmune Disease. Front Immunol. 2019 June 11;10:1090.

4. Lamberts A, Euverman HI, Terra JB, Jonkman MF, Horváth B. Effectiveness and safety of rituximab in recalcitrant pemphigoid diseases. Front Immunol. 2018;9(FEB):1–9.

5. Koga H, Prost-Squarcioni C, Iwata H, Jonkman MF, Ludwig RJ, Bieber K. Epidermolysis Bullosa Acquisita: The 2019 Update. Front Med [Internet]. 2019;5(January). Available from: https://www.frontiersin.org/article/10.3389/fmed.2018.00362/full

6. Bieber K, Witte M, Sun S, Hundt JE, Kalies K, Dräger S, et al. T cells mediate autoantibody-induced cutaneous inflammation and blistering in epidermolysis bullosa acquisita. Sci Rep. 2016;6(December):1–13.

7. Hübner F, Recke A, Zillikens D, Linder R, Schmidt E. Prevalence and Age Distribution of Pemphigus and Pemphigoid Diseases in Germany. J Invest Dermatol. 2016;136(12):2495– 8.

8. Joly P, Baricault S, Sparsa A, Bernard P, Bédane C, Duvert-Lehembre S, et al. Incidence and mortality of bullous pemphigoid in France. J Invest Dermatol. 2012;132(8):1998–2004.

9. Kridin K, Bergman R. Mortality in Patients with Bullous Pemphigoid: A Retrospective Cohort Study, Systematic Review and Meta-analysis. Acta Derm Venereol. 2018;0.

10. Polansky M, Eisenstadt R, DeGrazia T, Zhao X, Liu Y, Feldman R. Rituximab therapy in patients with bullous pemphigoid: A retrospective study of 20 patients. J Am Acad Dermatol. 2019 July 1;81(1):179–86.

11. Ujiie H, Rosmarin D, Schön MP, Ständer S, Boch K, Metz M, et al. Unmet Medical Needs in Chronic, Non-communicable Inflammatory Skin Diseases. Front Med. 2022;9:875492.

12. Boch K, Zirpel H, Thaci D, Mruwat N, Zillikens D, Ludwig RJ, et al. Mortality in eight autoimmune bullous diseases: A global large-scale retrospective cohort study. J Eur Acad Dermatol Venereol JEADV. 2023 Apr;37(4):e535–7.

13. Kridin K, Bieber K, Vorobyev A, Moderegger EL, Hernandez G, Schmidt E, et al. Risk of death, major adverse cardiac events and relapse in patients with bullous pemphigoid treated with systemic or topical corticosteroids. Br J Dermatol. 2024 Oct 1;191(4):539–47.

14. Goebeler ME, Bargou RC. T cell-engaging therapies — BiTEs and beyond. Nat Rev Clin Oncol. 2020 July;17(7):418–34.

15. Ellebrecht CT, Bhoj VG, Nace A, Choi EJ, Mao X, Cho MJ, et al. Reengineering chimeric antigen receptor T cells for targeted therapy of autoimmune disease. Science. 2016 July 8;353(6295):179–84.

16. Lee J, Lundgren DK, Mao X, Manfredo-Vieira S, Nunez-Cruz S, Williams EF, et al. Antigen-specific B cell depletion for precision therapy of mucosal pemphigus vulgaris. J Clin Invest. 2020 Oct 26;130(12):6317–24.

17. Chang DJ, Basu S, Micheletti R, Maverakis E, Marinkovich M, Porter DL, et al. LB952 A phase 1 trial of DSG3-CAART cells in mucosal-dominant pemphigus vulgaris (mPV) patients: Preliminary data. J Invest Dermatol. 2022 Aug;142(8):B18.

18. Cliff ERS, Kelkar AH, Russler-Germain DA, Tessema FA, Raymakers AJN, Feldman WB, et al. High Cost of Chimeric Antigen Receptor T-Cells: Challenges and Solutions. Am Soc Clin Oncol Educ Book. 2023 June;(43):e397912.

19. Braun T, Kuschel F, Reiche K, Merz M, Herling M. Emerging T-cell lymphomas after CAR T-cell therapy. Leukemia [Internet]. 2025 Apr 7 [cited 2025 May 9]; Available from: https://www.nature.com/articles/s41375-025-02574-x

20. Subklewe M, Magno G, Gebhardt C, Bücklein V, Szelinski F, Arévalo HJR, et al. Application of blinatumomab, a bispecific anti-CD3/CD19 T-cell engager, in treating severe systemic sclerosis: A case study. Eur J Cancer. 2024 June;204:114071.

21. Alexander T, Krönke J, Cheng Q, Keller U, Krönke G. Teclistamab-Induced Remission in Refractory Systemic Lupus Erythematosus. N Engl J Med. 2024 Sept 5;391(9):864–6.

22. Bucci L, Hagen M, Rothe T, Raimondo MG, Fagni F, Tur C, et al. Bispecific T cell engager therapy for refractory rheumatoid arthritis. Nat Med. 2024 June;30(6):1593–601.

23. Perico L, Casiraghi F, Sônego F, Todeschini M, Corna D, Cerullo D, et al. Bi-specific autoantigen-T cell engagers as targeted immunotherapy for autoreactive B cell depletion in autoimmune diseases. Front Immunol. 2024 Feb 26;15:1335998.

24. Hagen M, Bucci L, Böltz S, Nöthling DM, Rothe T, Anoshkin K, et al. BCMA-Targeted T-Cell– Engager Therapy for Autoimmune Disease. N Engl J Med. 2024 Sept 5;391(9):867–9.

25. Moreau P, Garfall AL, Van De Donk NWCJ, Nahi H, San-Miguel JF, Oriol A, et al. Teclistamab in Relapsed or Refractory Multiple Myeloma. N Engl J Med. 2022 Aug 11;387(6):495–505.

26. Kantarjian H, Stein A, Gökbuget N, Fielding AK, Schuh AC, Ribera JM, et al. Blinatumomab versus Chemotherapy for Advanced Acute Lymphoblastic Leukemia. N Engl J Med. 2017 Mar 2;376(9):836–47.

27. Bangolo A, Amoozgar B, Mansour C, Zhang L, Gill S, Ip A, et al. Comprehensive Review of Early and Late Toxicities in CAR T-Cell Therapy and Bispecific Antibody Treatments for Hematologic Malignancies. Cancers. 2025 Jan 17;17(2):282.

28. Kim BR, Lee KH, Paik K, Kim M, Bae JM, Choi CW, et al. Comorbid diseases in bullous pemphigoid: A population-based case–control study. J Dermatol. 2025 Mar;52(3):460–71.

29. Olbrich H, Patzelt S, Gaffal E, Kridin K, Curman P, Schmidt E, et al. Risk of oesophageal strictures in mucous membrane pemphigoid: Insights from a real-world cohort study. J Eur Acad Dermatol Venereol. 2025 Feb 26;jdv.20603.

30. Tigges M, Dräger S, Piccini I, Bieber K, Vorobyev A, Edelkamp J, et al. Pemphigoid disease model systems for clinical translation. Front Immunol. 2025 Mar 17;16:1537428.

31. Iwata H, Bieber K, Tiburzy B, Chrobok N, Kalies K, Shimizu A, et al. B cells, dendritic cells, and macrophages are required to induce an autoreactive CD4 helper T cell response in experimental epidermolysis bullosa acquisita. J Immunol Baltim Md 1950. 2013;191(6):2978–88.

32. Ghorbanalipoor S, Emtenani S, Parker M, Kamaguchi M, Osterloh C, Pigors M, et al. Cutaneous kinase activity correlates with treatment outcomes following PI3K delta inhibition in mice with experimental pemphigoid diseases. Front Immunol. 2022 Sept 28;13:865241.

33. Kasprick A, Hofrichter M, Smith B, Ward P, Bieber K, Shock A, et al. Treatment with anti-neonatal Fc receptor (FcRn) antibody ameliorates experimental epidermolysis bullosa acquisita in mice. Br J Pharmacol. 2020;177(10):2381–92.

34. Müller S, Behnen M, Bieber K, Möller S, Hellberg L, Witte M, et al. Dimethylfumarate Impairs Neutrophil Functions. J Invest Dermatol. 2016 Jan;136(1):117–26.

35. Covell DG, Barbet J, Holton OD, Black CD, Parker RJ, Weinstein JN. Pharmacokinetics of monoclonal immunoglobulin G1, F(ab’)2, and Fab’ in mice. Cancer Res. 1986 Aug;46(8):3969–78.

36. Benjamin JE, Stein AS. The role of blinatumomab in patients with relapsed/refractory acute lymphoblastic leukemia. Ther Adv Hematol. 2016 June;7(3):142–56.

37. Mohan N, Ayinde S, Peng H, Dutta S, Shen Y, Falkowski VM, et al. Structural and functional characterization of IgG- and non-IgG-based T-cell-engaging bispecific antibodies. Front Immunol. 2024 May 28;15:1376096.

38. Bumma N, Richter J, Jagannath S, Lee HC, Hoffman JE, Suvannasankha A, et al. Linvoseltamab for Treatment of Relapsed/Refractory Multiple Myeloma. J Clin Oncol. 2024 Aug;42(22):2702–12.

39. Rafidi H, Rajan S, Urban K, Shatz-Binder W, Hui K, Ferl GZ, et al. Effect of molecular size on interstitial pharmacokinetics and tissue catabolism of antibodies. mAbs. 2022 Dec 31;14(1):2085535.

40. Bhoj VG, Arhontoulis D, Wertheim G, Capobianchi J, Callahan CA, Ellebrecht CT, et al. Persistence of long-lived plasma cells and humoral immunity in individuals responding to CD19-directed CAR T-cell therapy. Blood. 2016 July 21;128(3):360–70.

41. O’Connor BP, Raman VS, Erickson LD, Cook WJ, Weaver LK, Ahonen C, et al. BCMA Is Essential for the Survival of Long-lived Bone Marrow Plasma Cells. J Exp Med. 2004 Jan 5;199(1):91–8.

42. Schroeder HW, Radbruch A, Berek C. B-Cell Development and Differentiation. Clin Immunol. 2019;107-118.e1.

43. Hammarlund E, Thomas A, Amanna IJ, Holden LA, Slayden OD, Park B, et al. Plasma cell survival in the absence of B cell memory. Nat Commun. 2017 Nov 24;8(1):1781.

44. Tedder TF. CD19: a promising B cell target for rheumatoid arthritis. Nat Rev Rheumatol. 2009 Oct;5(10):572–7.

45. Shi M, Wang J, Huang H, Liu D, Cheng H, Wang X, et al. Bispecific CAR T cell therapy targeting BCMA and CD19 in relapsed/refractory multiple myeloma: a phase I/II trial. Nat Commun. 2024 Apr 20;15(1):3371.

46. Leo O, Foo M, Sachs DH, Samelson LE, Bluestone JA. Identification of a monoclonal antibody specific for a murine T3 polypeptide. Proc Natl Acad Sci. 1987 Mar;84(5):1374–8.

47. Peipp M, Lammerts Van Bueren JJ, Schneider-Merck T, Bleeker WWK, Dechant M, Beyer T, et al. Antibody fucosylation differentially impacts cytotoxicity mediated by NK and PMN effector cells. Blood. 2008 Sept 15;112(6):2390–9.

48. Sitaru C, Chiriac MT, Mihai S, Büning J, Gebert A, Ishiko A, et al. Induction of complement-fixing autoantibodies against type VII collagen results in subepidermal blistering in mice. J Immunol Baltim Md 1950. 2006;177(5):3461–8.

49. Bieber K, Koga H, Nishie W. In vitro and in vivo models to investigate the pathomechanisms and novel treatments for pemphigoid diseases. Exp Dermatol. 2017;26(12):1163–70.

50. Gross N, Marketon J, Mousavi S, Kalies K, Ludwig RJ, Bieber K. Inhibition of interferon gamma impairs induction of experimental epidermolysis bullosa acquisita. Front Immunol. 2024 May 10;15:1343299.

51. Dräger S, Kalies K, Sidronio TB, Witte M, Ludwig RJ, Bieber K. Increased TREM-1 expression in inflamed skin has no functional impact on the pathogenesis of cutaneous disorders. J Dermatol Sci. 2017 May 29;88:139–55.

52. Pipi E, Kasprick A, Iwata H, Goletz S, Hundt JE, Sadeghi H, et al. Multiple Modes of Action Mediate the Therapeutic Effect of Intravenous IgG in Experimental Epidermolysis Bullosa Acquisita. J Invest Dermatol. 2022;142(6):1552–1564.e8.

53. Hammers CM, Bieber K, Kalies K, Banczyk D, Ellebrecht CT, Ibrahim SM, et al. Complement-Fixing Anti-Type VII Collagen Antibodies Are Induced in Th1-Polarized Lymph Nodes of Epidermolysis Bullosa Acquisita-Susceptible Mice. J Immunol. 2011 Nov 15;187(10):5043– 50.

