## Supplemental information for "CD19×CD3 bispecific T cell engager treatment induces remission in experimental pemphigoid disease"

**Supplemental Material**

**Table S1: Weight and total burden of mice in dosing and route application finding experiments.** Weight (g) and total burden of each mouse were assessed in each group at the start and the end of the experiment. Here, values are depicted for the route application and dosage finding experiments as mean with standard deviation. No loss in weight and no or only mild total burden could be observed with “0” as “no burden”, “1-10” as “mild burden”, “11-20” as “mid burden”, and “>20” as “severe burden” following immediate stop of the experiment and sacrificing of the individual mouse.

|  |  |  | **0.1 µg** | **1 µg** | **10 µg** | **25 µg** |
| --- | --- | --- | --- | --- | --- | --- |
| **i.p.** | weight (g) | start | 22.67 ± 4.73 | 26.00 ± 6.08 | 29.00 ± 2.00 | 23.67 ± 4.04 |
|  |  | end | 23.33 ± 4.04 | 26.33 ± 4.73 | 29.00 ± 1.73 | 23.67 ± 3.79 |
|  | total burden | start | 0.00 ± 0.00 | 0.00 ± 0.00 | 0.00 ± 0.00 | 0.00 ± 0.00 |
|  |  | end | 0.00 ± 0.00 | 0.34 ± 0.58 | 0.34 ± 0.58 | 0.34 ± 0.58 |
| **i.v.** | weight (g) | start | 27.33 ± 2.08 | 28,67 ± 2.08 | 22.33 ± 5.77 | 25.00 ± 5.29 |
|  |  | end | 28.00 ± 2.00 | 27.33 ± 2.52 | 23.33 ± 4.93 | 25.00 ± 5.29 |
|  | total burden | start | 0.00 ± 0.00 | 0.00 ± 0.00 | 0.00 ± 0.00 | 0.00 ± 0.00 |
|  |  | end | 0.00 ± 0.00 | 2.34 ± 2.31 | 0.00 ± 0.00 | 0.00 ± 0.00 |

**Table S2: Comparison of change in weight and total burden of mice after one time or multiple injections of CD19xCD3 TCE.** Weight (g) and total burden of each mouse were assessed in each group at the start and the end of the experiment. Here, values are depicted for the one-time injection of 1 µg CD19xCD3 TCE i.p. as compared to 2-3x injections as mean with standard deviation. No loss in weight and no or only mild total burden could be observed with “0” as “no burden”, “1-10” as “mild burden”, “11-20” as “mid burden”, and “>20” as “severe burden” following immediate stop of the experiment and sacrificing of the individual mouse. TCE: T cell engager.

|  |  |  | **96 h** | **168 h** |
| --- | --- | --- | --- | --- |
| **1x injection** | weight (g) | start | 31.67 ± 5.77 | 26.00 ± 6.08 |
|  |  | end | 30.67 ± 5.77 | 26.33 ± 4.73 |
|  | total burden | start | 0.00 ± 0.00 | 0.00 ± 0.00 |
|  |  | end | 1.00 ± 1.00 | 0.00 ± 0.00 |
| **2-3x injection** | weight (g) | start | 26.00 ± 6.00 | 24.67 ± 2.89 |
|  |  | end | 27.00 ± 4.58 | 24.33 ± 3.21 |
|  | total burden | start | 0.00 ± 0.00 | 0.00 ± 0.00 |
|  |  | end | 0.34 ± 0.58 | 0.34 ± 0.58 |

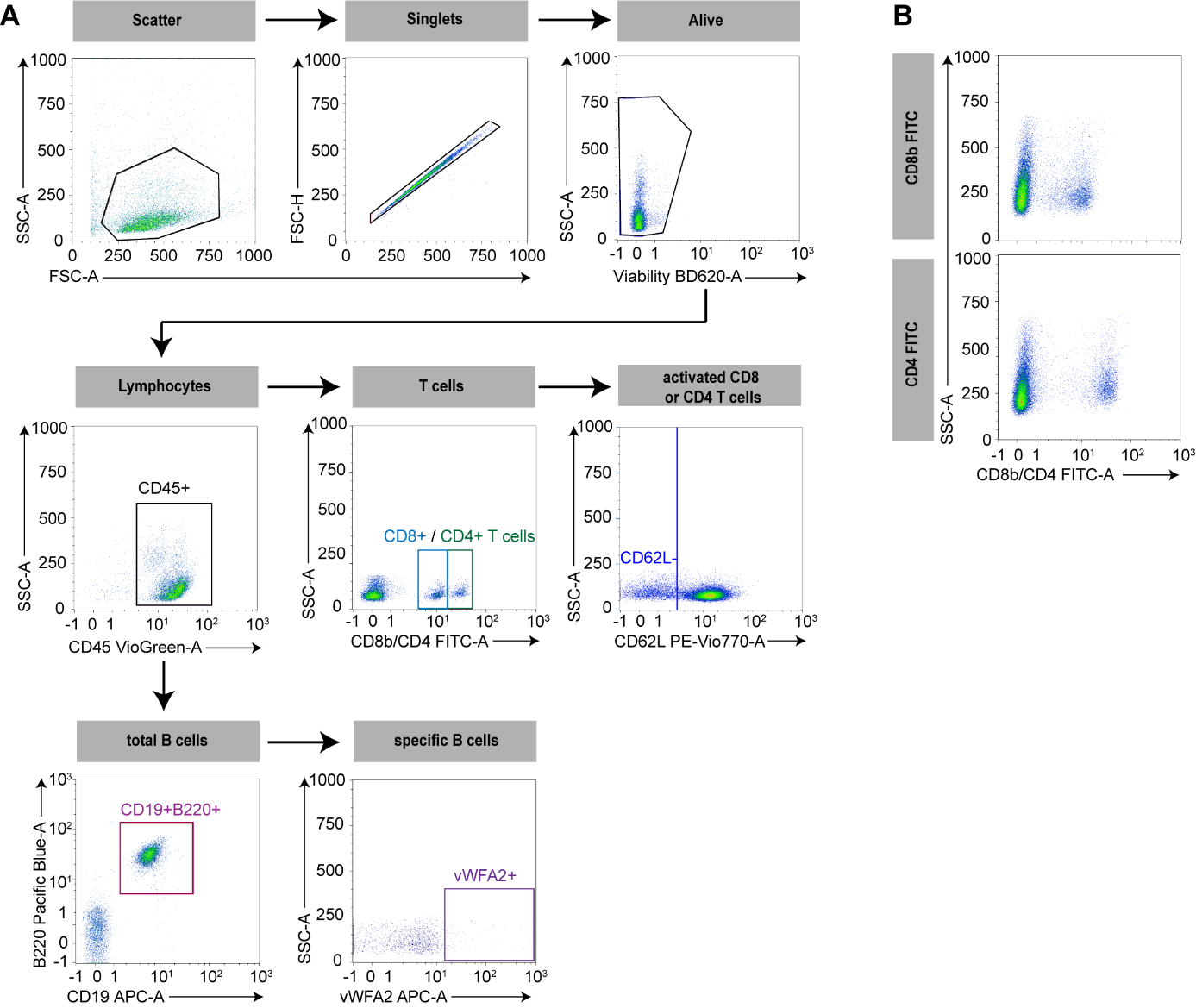

**Figure S1: Gating strategy for the detection of B and T cells by flow cytometry.** **(A)** Cells were gated for leukocytes (FSC-A and SSC-A) and singlets (FSC-H compared with FSC-A). Single cells were then differentiated between alive and dead. Only living cells were further gated on their positive CD45 expression and then the positive B cell (B220 and CD19) and T cell (CD4 and CD8) population. B220^+^CD19^+^ B cells were further analyzed if they were specific for COL7^vWFA2^ (vWFA2+). CD4^+^ and CD8^+^ T cells were both gated for their expression of CD62L and the negative population represented the activated CD8 or CD4 T cell population. **(B)** Single stainings of the antibodies anti-CD8 FITC and anti-CD4 FITC and the corresponding mean fluorescence signal in the FITC channel.

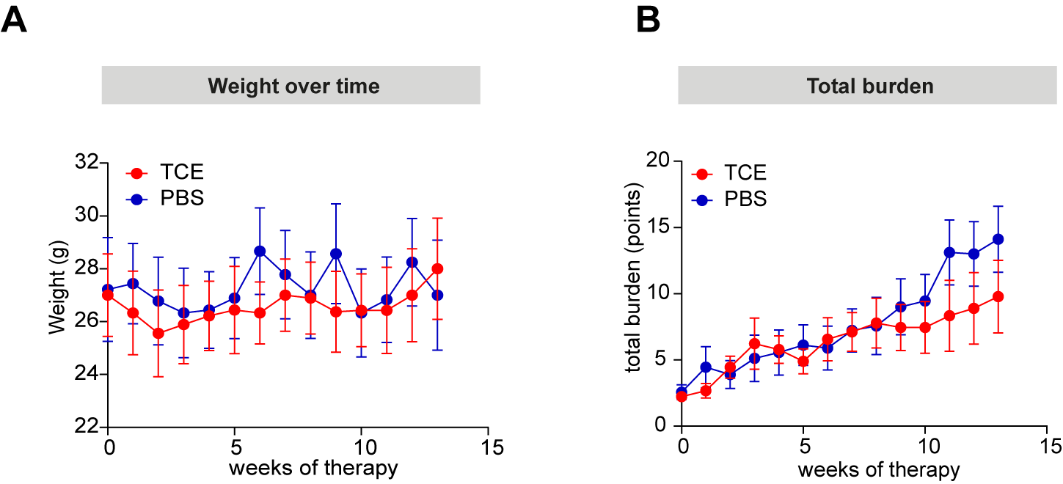

**Figure S2: Similar development of weight change and total burden of treated or not-treated mice after randomization.** Comparison of **(A)** weight (g) and **(B)** total burden of immunized mice of treated (TCE) and control (PBS) group after randomization until week 13 of therapy. No difference could be observed between the two groups. Data are presented as mean values ± SD. TCE: T cell engager.

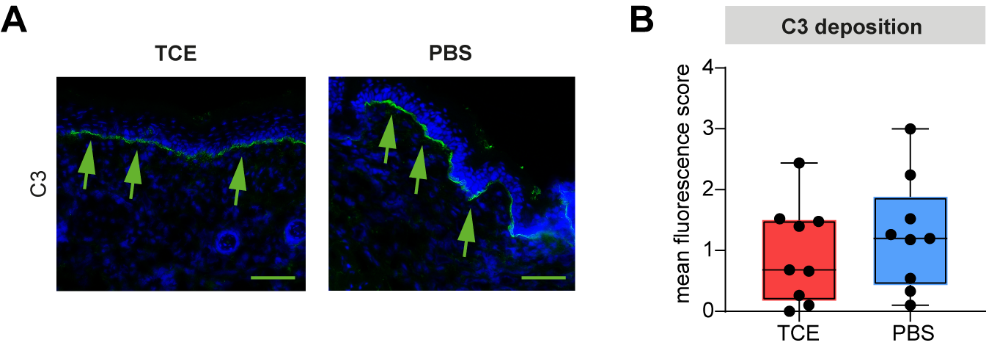

**Figure S3: No difference in C3 deposition in CD19xCD3 TCE treated mice to control mice in the immunized mouse model of epidermolysis bullosa acquisita. (A)** Representative pictures of the direct immunofluorescence staining for C3 deposition at the dermal-epidermal junction (green arrows) for both groups. **(B)** Analysis showed no difference between the two groups. Data are presented as medians (black line), 25^th^/75^th^ percentiles (boxes), and max/min values (error bars). Scale bar: 100 µm. TCE: T cell engager.
